# Determinants of Self-referral among Outpatients at Referral Hospitals in East Wollega, Western Ethiopia

**DOI:** 10.1101/540476

**Authors:** Edosa Tesfaye Geta*, Yibeltal Siraneh Belete, Elias Ali Yesuf

## 1. Introduction

Patient self-referral is a condition when patients refer themselves to higher level health facilities without having to see anyone else first or without being told to refer themselves by another health professional [1]. Primary health care facilities need to maintain a close relationship between all the levels of a health system and the referral linkage of health facilities is very important in providing health care for the people of any country [2].

In many developing countries, a high proportion of clients or patients seen at the outpatient departments at secondary health care facilities could be appropriately looked after at primary health care centers [3].

Ethiopia has successfully implemented its strategy of expanding and rehabilitating primary health care facilities to make health services closer to the community and easily accessible and to ensure further decentralization efforts have been done. However, the health care utilization at nearest primary health care facilities in the country is still low [4].

Referral functions are usually invoked to justify the privileged resource allocations of higher health facilities. To counter-argue with hard evidence of reduced referral caseloads may strengthen the bargaining position of lower-level facilities and the expansion of overcrowded referral health facilities in fact attending an excessive proportion of common conditions without offering significant comparative benefits to these patients can be discouraged [5].

According to the study conducted in South Africa, more than 50% of the patients seen at referral health facilities outpatient departments could have been managed at the primary level health facilities. Most acute cases at the hospital were self-referral where the patients probably did not seek health care until they became severely ill or sought care at other levels of health care, and then came to the referral hospital [6].

In spite of primary health care elements providing essential care and prevention resources for rural populations in Ethiopia, patients often access secondary levels of care without first using primary health care provided by primary level health facilities [7]. The study conducted in western Ethiopia at general hospital revealed 82% of patients bypassed the first level referral facilities and 74.9% of them did not first contact nearest health facilities [8] and the fundamental message of referral system is still unchanged [9].

Referral flows should be understood because of their efficiency and effectiveness implications. Little is known about determinants of outpatient self-referral in Ethiopia. So determining the magnitude and identifying the factors that contribute to self-referral to referral health facilities are very important in provision of health care in the country.

## 2. Methods and Participants

### Study Design and Participants

This study was conducted from December 01 to 30, 2017 using a cross-sectional facility based study design in East Wollega zone of Oromia National Regional state, Ethiopia.

All patients visited the outpatient departments of the study health facilities willing to participate in this study were included in the study after taking consent and for patient whose age was less than 15year; their care givers/parents were interviewed and patients with serious physical, mental problems and emergency cases were excluded from the final interview of the study. Patients whose age was less than 15 year and without care giver/parents were excluded.

### Sampling and Data Collection Procedures

The required sample size for the study was determined by using formula of single population proportion [10] with the assumption of 50% outpatient self-referral rate, 95% confidence interval with 5% margin of error and 5% non-response rate;

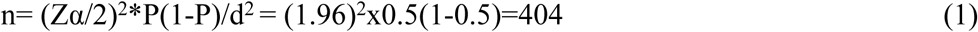

Data was collected from outpatients visited Nekemte referral hospital. Nekemte referral hospital was selected purposely because the other referral hospital in east Wollega was less than one year since it was established and became functional during the study period. By using systematic sampling method 404 respondents were selected. The interval of the respondents for the interview was determined by dividing the daily average total number of outpatient flow excluding emergency outpatient departments by the number of participants planned to be interviewed daily. Therefore, every K^th^ from Nekemte referral hospital (180/24=7th) and from registration books in triage, simple random sampling technique was employed to select the first participant.

The structured questionnaire was developed in English after reviewing relevant literatures and the questionnaire was translated in to local language Afan Oromo wich was used for interview. Three data collectors and one supervisor were assigned and trained for the purpose data collection. Both supervisor and data collectors those had BSc degree and experience were assigned.

Data was collected by face-to-face interview using structured questionnaire. Patients who completed their outpatient services and return to leave the health facility and willing to participate in this study were interviewed (exit interview).

### Study Variables

The dependent variable is outpatient self-referral and interdependent variables are socio-demographic characteristics, location of health care facilities, access to public transport, proximity to home of health care facilities, proximity to work place of health facility, affordability of health services, health services provider type, type of proximal health facility, level of health facility, length of waiting time in health facility, reputation of health facility, availability of credit facility, availability of adequate drugs, availability of advanced laboratory service, availability of radiological services, perceived quality of services, perceived severity of illness, information about referral system, self-reported chief complaints, health care provider courtesy and respect.

### Data Management and Analysis

The completed questionnaires were being checked for completeness, consistency and coded by the principal investigator and supervisors. Questionnaires for participants were translated into local language (Afan Oromo) and retranslated back into English to ensure its consistency. The questionnaires were pre-tested using 40(5%) of sample size at Wollega University referral hospital and to ensure that whether it is clear or not for respondents.

After completeness of each questionnaires checked, data entry and analysis were made using SPSS for windows version 20 software. Descriptive statistics of frequency was performed for all study variables. Simple logistic regression was performed to identify association between each independent and dependent variables.

All variables associated with outpatient self-referral bypassing primary health facilities in bivariate analysis were considered as candidate variable with p≤0.25. Those candidate variables were entered into multivariable logistic regressions to identify predictors of self-referral at p<0.05 to declare level of significance.

## 3. Results

A total of 404 of outpatients were included in the study making 391(96.8%) of response rate. Among 391 outpatients visited Nekemte referral hospital outpatient departments 330(84.4%) were self-referred outpatients who had bypassed the nearest health facilities. Among self-referred outpatients, more than half of patients 177(53.6%) were females and majority of the self-referred outpatients 212(64.2%) were from urban and 212(64.3%) were literate (***Table 1***).

**Table 1:** Socio-demographic characteristics and self-referral of outpatients visited outpatient departments at Nekemte referral hospital in East Wollega, Ethiopia 2017.

| Socio-demographic characteristics |  | Self-referral |  |  |
| --- | --- | --- | --- | --- |
|  |  | Yes<br>N=330 (%) | No<br>N=61(%) | Total<br>N=391 |
| Sex | Male | 153(46.4) | 19(31.1) | 172 |
|  | Female | 177(53.6) | 42(26.2) | 219 |
| Marital status | Single | 146(44.2) | 16(41.0) | 162 |
|  | Married | 184(55.8) | 45(73.8) | 229 |
| Place of Residence | Urban | 212(64.2) | 26(42.6) | 238 |
|  | Rural | 118(35.8) | 35(57.4) | 153 |
| Educational status | Illiterate | 106(32.1) | 29(47.5) | 135 |
|  | Read & Write | 12(3.6) | 3(4.9) | 15 |
|  | 1<sup>st</sup> cycle (grade 1-4) | 45(13.6) | 1(1.6) | 46 |
|  | 2<sup>nd</sup> cycle(grade 5-8) | 51(15.5) | 9(14.8) | 60 |
|  | Secondary & Prep(9-12) | 50(15.2) | 6(9.8) | 56 |
|  | Diploma | 40(12.1) | 10(16.4) | 50 |
|  | Degree and above | 26(7.9) | 3(4.9) | 29 |
| Employment status | Employed | 69(20.9) | 19(31.1) | 88 |
|  | Unemployed | 261(79.1) | 42(68.9) | 303 |

### Reasons for Outpatients Self-referral to Referral hospitals

From a total of 330 self-referred outpatients interviewed at Nekemte referral hospital 301(91.2%) of patients responded as the study hospital was not the health facility closest to their residence place and out of 301; 160(53.2%) did not visit the nearest health facilities for their current health problems. The type of proximal health facilities bypassed 46(15.5%) were public hospitals, 145(48.8%) were public health centers and 110(35.7%) were private clinics.

Among the respondents 190(48.6%) visited Nekemte referral hospital for their first time and 201(51.4%) had previous experience of visiting the facility and the majority of self-referred outpatients 239(72.4%) had no referral information of health facilities referral linkage. The perceived severity of illness of self-referred patients, 124(37.6%) were severe and 206(62.4%) were mild whereas the majority of category of perceived self-reported chief complaints 250(75.8%) were acute.

The interviewed outpatients reasoned out different reasons from their previous experience of nearest health facilities visits why they preferred to visit outpatient departments of referral hospital bypassing the proximal health facilities that they visited in previous.

From a total of 330 self-referred out patients for 169(51.2%) participants, the proximal health facility location was not convenient and 259(78.5%) of the proximal health facilities had no transportation access to be visited and 239 (72.4%) participants were not confident to get health care providers type they want to see for their health problems at the proximal facility.

From their previous visits of proximal health facilities, 153(46.4%) of participants said the health care providers courtesy and respect at the facility was not good and 289 (87.6%) preferred medical doctors to other clinicians to see firstly for their any health problems and visited the hospital, 222(67.3%) preferred to visit referral hospital to get more and detail explanation about their health problems that they did not get during proximal health facility visit.

The other reasons for self-referral of outpatients to referral hospital were the expectation of availability of adequate drugs and laboratory services at the hospital. From self-referred outpatients; 194(58.8%) and 216(65.5%) participants did not get adequate drugs and laboratory services respectively as they want at proximal health facilities during their previous visits (***Table 2***).

**Table 2:** Reasons for outpatient self-referral to Nekemte referral hospital in East Wollega, Ethiopia, 2017

| Reasons for Self-referral |  | Self-referral |  |  |
| --- | --- | --- | --- | --- |
|  |  | Yes<br>n=330 (%) | No<br>n=61(%) | Total<br>n=391 |
| Is NRH the proximal facility to your residence place? | Yes | 29(8.8) | 0 | 29 |
|  | No | 301(91.2) | 61(100) | 362 |
| Proximal health facility type | Hospital | 46(15.5) | 9(14.8) | 55 |
|  | Health center | 145(48.8) | 39(63.9) | 184 |
|  | Private clinic | 110(35.7) | 13(21.3) | 123 |
| Referral Information | Yes | 91(27.6) | 30(49.2) | 121 |
|  | No | 239(72.4) | 31(50.8) | 270 |
| Visiting of nearest HF for current health problem | Yes | 141(46.8) | 43(70.5) | 184 |
|  | No | 160(53.2) | 18(29.5) | 178 |
| Visiting the proximal HF in last 12 months | Yes | 292(97) | 60(98.4) | 352 |
|  | No | 9(3) | 1(1.6) | 10 |
| How often you visit NRH? | First time | 154(46.7) | 36(59) | 190 |
|  | Occasionally | 79(23.9) | 11(18.0) | 90 |
|  | Often | 97(29.4) | 14(23) | 111 |
| Perceived severity of illness | Severe | 124(37.6) | 13(21.3) | 137 |
|  | Mild | 206(62.4) | 48(78.7) | 254 |
| Self reported chief complaints | Chronic | 80(24.2) | 23(37.7) | 103 |
|  | Acute | 250(75.8) | 38(62.3) | 288 |
| Access to public transport to visit the proximal HF | Yes | 71(21.5) | 30(49.2) | 101 |
|  | No | 259(78.5) | 31(50.8) | 290 |
| Convenient location of the proximal HF | Yes | 161(48.8) | 40(65.6) | 201 |
|  | No | 169(51.2) | 21(34.4) | 190 |
| Affordability of Services from the proximal HF | Yes | 275 (83.3) | 47 (77) | 322 |
|  | No | 55 (16.7) | 14 (23) | 69 |
| Confident to get the providers from proximal HF? | Yes | 91(27.6) | 36(59) | 127 |
|  | No | 239(72.4) | 25(41) | 264 |
| Patients' preference of health care provider to see firstly | Doctors | 289(87.6) | 58(95.1) | 347 |
|  | BSc/Ho | 41(12.4) | 3(4.9) | 44 |
| Did you get explanation about your health problems | Yes | 108(32.7) | 26(42.7) | 134 |
|  | No | 222(67.3) | 53(57.3) | 275 |
| Was provider courtesy and respect good at proximal HF? | Yes | 153(46.4) | 37(60.7) | 190 |
|  | No | 177(53.6) | 24(39.3) | 201 |
| Time to get service short at the proximal HF? | Yes | 255(77.3) | 41(67.2) | 296 |
|  | No | 75(22.7) | 20(32.8) | 95 |
| Was the reputation of the proximal HF good? | Yes | 128(38.8) | 25(41) | 153 |
|  | No | 202(61.2) | 36(59) | 238 |
| Did you get adequate drugs in the HF during your visit? | Yes | 136(41.2) | 50(82) | 186 |
|  | No | 194(58.8) | 11(18) | 205 |
| Did you get laboratory services at proximal HF? | Yes | 114(34.5) | 44(72.1) | 158 |
|  | No | 216(65.5) | 17(27.9) | 233 |
| Availability of radiology services in the Proximal HF? | Yes | 35(10.6) | 14(23%) | 49 |
|  | No | 295(89.4) | 47(77%) | 342 |
| Rate the quality of services at the proximal HF | Good | 113(34.2) | 37(60.7) | 150 |
|  | Poor | 217(65.8) | 24(39.3) | 241 |

**Notes:** NRH=Nekemte Referral Hospital, HF= Health Facility, HO= Health Officer

### Predictors of Outpatient Self-referrals to Referral hospitals

From multiple logistic regression analysis (***Table 3***), the referral information on linkage of health facilities was very strongly associated with outpatient self-referral to visit outpatient department of referral hospitals (AOR and 95% CI**=**0.324(0.150-0.696). That means outpatients who had information about referral linkage of health facilities were 68% less likely to self-refer to referral hospitals than those patients that had no any information about the referral linkage of health facilities.

**Table 3:** Factors influencing Outpatient self-referral to Nekemte referral hospital in East Wollega, Oromia National Regional state, Ethiopia, 2017

| Variables | Self-referral |  | OR and 95%CI |  |
| --- | --- | --- | --- | --- |
|  | Yes (%) | No (%) | COR | AOR |
| **Sex** | 330 | 61 |  |  |
| Male | 153(46.4) | 19(31.1) | 1.911(1.066-3.425) | 1.807 (0.827-3.948) |
| Female | 177(53.6) | 42(68.9) | 1 | 1 |
| **Marital Status** | 330 | 61 |  |  |
| Single | 146(44.2) | 16(41.0) | 2.232(1.212-4.109) | 1.252 (0.492-3.189) |
| Married | 184(55.8) | 45(73.8) | 1 | 1 |
| **Residence place** | 330 | 61 |  |  |
| Urban | 212(64.2) | 26(42.6) | 2.419(1.388-4.21) | 1.978 (0.927-4.222) |
| Rural | 118(35.8) | 35(57.4) | 1 | 1 |
| **Educational status** | 330 | 61 |  |  |
| Illiterate | 106(32.1) | 29(47.5) | 0.422(.119-1.492) | 0.193 (0.028-1.327) |
| Read & Write | 12(3.6) | 3(4.9) | 0.462(0.081-2.630) | 0.391 (0.024-6.346) |
| 1<sup>st</sup> cycle (1-4) | 45(13.6) | 1(1.6) | 5.192(.513-52.524) | 1.377 (0.081-23.260) |
| 2<sup>nd</sup> cycle(5-8) | 51(15.5) | 9(14.8) | 0.654(0.163-2.623) | 0.287 (0.037-2.247) |
| Grade (9-12) | 50(15.2) | 6(9.8) | 0.962(.222-4.160) | 0.365 (0.056-2.946) |
| Diploma | 40(12.1) | 10(16.4) | 0.462(0.116-1.837) | 0.463 (.063-3.412) |
| Degree & above | 26(7.9) | 3(4.9) | 1 | 1 |
| **Jobs status** | 330 | 61 |  |  |
| Employed | 69(20.) | 19(31.1) | 0.584(0.320-1.068) | 0.567 (0.250-1.286) |
| Unemployed | 261(79.1) | 42(68.9) | 1 | 1 |
| **Proximal HF type** | 298 | 61 |  |  |
| Hospital | 46(15.5) | 9(14.8) | 0.627(0.250-1.569) | 0.731 (0.150-3.557) |
| Health center | 145(48.8) | 39(63.9) | 0.456(0.232-0.896) | 0.716 (.236-2.176) |
| Private clinic | 110(35.7) | 13(21.3) | 1 | 1 |
| **Current visiting of nearest HF** | 301 | 61 |  |  |
| Yes | 141(46.8) | 43(70.5) | 0.369(0.203-0.669) | 0.478 (0.224-1.019) |
| No | 160(53.2) | 18(29.5) | 1 | 1 |
| **Referral linkage information** | 330 | 61 |  |  |
| Yes | 91(27.6) | 30(49.2) | 0.393(0.225-0.687) | 0.324 (0.150-0.696)** |
| No | 239(72.4) | 31(50.8) | 1 | 1 |
| **Frequency of NRH visiting** | 330 | 61 |  |  |
| First time | 154(46.7) | 36(59) | 0.617(.317-1.204) | 0.768 (0.296-1.992) |
| Occasionally | 79(23.9) | 11(18) | 1.037(0.446-2.410) | 1.363 (0.433-4.286) |
| Often | 97(29.4) | 14(23) | 1 | 1 |
| **Perceived severity of illness** | 330 | 61 |  |  |
| Severe | 124(37.6) | 13(21.3) | 2.223(1.158-4.226) | 3.496 (1.473-8.297)** |
| Mild | 206(62.4) | 48(78.7) | 1 | 1 |
| **Self-reported chief complaint** | 330 | 61 |  |  |
| Chronic | 80(24.2) | 23(37.7) | 0.529(.297-0.940) | 0.903 (0.335-2.432) |
| Acute | 250(75.8) | 38(62.3) | 1 | 1 |
| **Affordability of Services from HF** | 330 | 61 |  |  |
| Yes | 275 (83.3) | 47 (77) | 1.489(0.767-2.891) | 0.459 (0.174-1.211) |
| No | 55 (16.7) | 14 (23) |  | 1 |
| **Convenience location of the HF** | 330 | 61 |  |  |
| Yes | 161(48.8) | 40(65.6) | 1 | 1 |
| No | 169(51.2) | 21(34.4) | 1.999 (1.130-3.538) | 3.750(1.560-9.017)** |
| **Access to transportation to visit HF** | 330 | 61 |  |  |
| Yes | 71(21.5) | 30(49.2) | 1 | 1 |
| No | 259(78.5) | 31(50.8) | 4.138 (2.350-7.288) | 3.879 (1.651-9.114)** |
| **Confident to get providers at HF** | 330 | 61 |  |  |
| Yes | 91 (27.6) | 36(59%) | 1 | 1 |
| No | 239 (72.4) | 25(41) | 3.782 (2.151-6.651) | 3.595 (1.554-8.317)** |
| **Patients' preference of provider type** | 330 | 61 |  |  |
| BSc/Ho | 41(12.4) | 3(4.9) | 0.655 (0.375-1.143) | 0.793 (0.172-3.646) |
| Doctors | 289(87.6) | 58(95.1) | 1 | 1 |
| **Did you get explanation at the HF?** | 330 | 61 |  |  |
| Yes | 108(32.7) | 26(42.7) | 0.655(0.375-1.143) | 0.926 (0.349-2.456) |
| No | 222(67.3) | 53(57.3) | 1 | 1 |
| **Care provider courtesy and respect** | 330 | 61 |  |  |
| Good | 153(46.4) | 153(46.4) | 0.561 (0.321-0.979) | 1.433 (0.612-3.359) |
| Poor | 177(53.6) | 177(53.6) | 1 | 1 |
| **Time to get service was short at t HF** | 330 | 61 |  |  |
| Yes | 255(77.3) | 41(67.2) | 1.659 (0.916-3.002) | 0.813(0.294-2.250) |
| No | 75(22.7) | 20(32.8) | 1 | 1 |
| **Availability of laboratory services** | 330 | 61 |  |  |
| Yes | 114 (34.5) | 44 (72.1) | 1 | 1 |
| No | 216 (65.5) | 17(27.9) | 4.904 (2.681-8.971) | 4.966 (2.199-11.216)*** |
| **Availability of drugs in the facility** | 330 | 61 |  |  |
| Yes | 136 (41.2) | 50(82) | 1 | 1 |
| No | 194 (58.8) | 11(18) | 6.484 (3.257-12.90) | 2.366 (1.013-5.526)* |
| **Did you get radiology services at HF** | 330 | 61 |  |  |
| Yes | 35(10.6) | 14(23%) | 2.511(1.257 | 0.893 (0.291-2.736) |
| No | 295(89.4) | 47(77%) | 1 | 1 |
| **Rate quality of services from the HF** | 330 | 61 |  |  |
| Good | 113(34.2) | 37(60.7) | 1 | 1 |
| Poor | 217(65.8) | 24(39.3) | 2.961(1.688-5.192) | 2.996(1.418-6.328)** |

**Notes:** \**P-value <0.05*, \*\**P-value <0.01*, \*\*\**P-value <0.001* and 1=reference, AOR = Adjusted Odd Ratio, COR = Crude Odd Ratio,
CI= Confidence Interval, NRH=Nekemte Referral Hospital, HF= Health Facility, HO= Health Officer

Regarding the perceived severity of illness, it was strongly associated with outpatient self-referral to referral hospitals (AOR and 95% CI=3.496(1.473-8.297). This odd ratio depicts that those patients perceived their illness as severe were 3.5 times more likely to self-refer to referral hospitals than those patients perceived their illness as mild.

The study result showed that, the confidence of patients to get health care providers they want at proximal health facilities was also strongly associated with outpatient self-referral to referral hospitals (AOR and 95% CI=3.595 (1.554-8.317). This indicates that patients those were not confident to get health care providers they want at proximal health facilities were 3.6 times more likely to self-refer to referral hospitals than those patients who were confident to get health care providers they want at proximal health facilities.

According to the study, the patients’ expectation about availability of advanced laboratory services and drugs at health facilities were significantly associated with outpatient self-referral to referral hospitals (AOR and 95% CI=4.966(2.199-11.216) and 2.366(1.013-5.526) respectively. This shows that the patients those did not expect to get laboratory services and drugs at proximal health facilities and had expectation of the availability of advanced laboratory and drugs at referral hospital almost five and 2.4 times more likely to self-refer to referral hospital respectively than those patients that obtained laboratory services and drugs at proximal health facilities.

The perceived quality of services was very strongly associated with outpatient self-referral to referral hospitals (AOR and 95% CI=2.996(1.418-6.328) and this indicates that odd of self-referral patients perceived quality of health services at proximal health facility was poor were almost three times more likely to self-refer to referral hospitals than those patients perceived the quality of health services at proximal health facility was good.

The convenience location and access to transportation of the health facilities were also strongly associated with outpatients self-referral to the referral hospitals (AOR of CI= 95%=3.750(1.560-9.017) and 3.879(1.651-9.114) respectively. This implies that the patients to whom proximal health facility location was inconvenient were 3.8 more likely to self-refer than those patients perceived the location proximal health facility was convenient and similarly those patients that had no access to transportation to visit proximal health facility almost four times more likely to self-refer than those had access to transportation to visit the nearest health facilities.

## 4. Discussion

According the study findings the proportion of outpatient self-referral to referral hospitals bypassing the proximal health facilities was 84.4%. This is finding shows the magnitude of patients’ self-referral was higher than the magnitude of patients’ self-referral in India which was 76.2% [11], in Tanzania 72.5% [12], but lower than that of Ghana which was 90% [13] and in Australia, where self-referral has been possible for over 30 years, approximately 65% of patients use self-referral [14].

This difference might be due to those studies findings show the proportion of general patients’ self-referral whereas this study shows the magnitude of outpatients’ self-referral to visit outpatient departments of referral hospitals.

According the study, the odd ratio depicts that those outpatients perceived their illness as severe were 3.5 more likely to self-refer to referral hospitals than those patients perceived their illness as mild (AOR of 95% CI=3.496 (1.473-8.297), but patients those had health facilities referral linkage information were 68% less likely to self-refer to referral hospitals than those patients that has no information about the referral linkage of health facilities (AOR of 95% CI=0.324 (0.150-0.696). The finding was consistent with the study findings conducted in Ethiopia those patients needing specialized care from higher level health facilities were less likely to have sought prior care at lower health facilities and in similar way patients that knew the closest health facility as first referral level were 76% less likely self-refer and those obtained information on referral systems were 35% less likely self-refer [7, 8].

Also other study in Kenya showed that respondent suffering from illness severity was found to be statistically significant to patient self-referral [15]. This might be when patients perceived their health problems as severe they may seek special health care from higher level health facilities and bypass the proximal health facility and when these patients obtained information on the types of health services and health care providers at proximal health facility and referral health facilities they might be able to utilize health services at proximal health facility.

Outpatients those were not confident to get all types of health care providers at proximal health facilities were 3.6 times more likely to self-refer to referral hospitals than those patients that were confident to get all types of health care providers at proximal health facilities (AOR of 95% CI=3.595(1.554-8.317). The study finding conducted in India showed that the factors of patients directly self-referral to secondary and tertiary referral health facilities are faith on the doctors and health facility 81% and availability of the specialists 54.5% [11] and the study conducted in Midwestern also showed that reasons for self-referral; preferences of health care providers type that was 37.5% of patients reported that they preferred to directly access a specialist to save time or to choose their own specialist [16].

This could be explained by the fact that from the finding of the study; 72.1% of self-referred patients were confident to get all types of health **c**are provider they want to see for their health problems at referral health facilities than the nearest health facilities and 87.6% of the self-referred patients preferred to see medical doctors firstly for their any health problems that were not available at primary level health facilities.

According to the study, 65.5% of the patients did not obtain laboratory services at proximal health facilities during their previous visit and preferred visiting referral hospitals to get advanced laboratory services. This shows that the patients that did not obtain laboratory services at proximal health facilities and expected the availability of laboratory services at referral hospitals almost five times more likely to self-refer to referral hospitals than those that obtained laboratory services at proximal primary health facilities (AOR of 95% CI=4.966(2.199-11.216). In addition to this the patients that did not obtain adequate drugs from proximal health facilities were 2.4 times more likely to self-refer to referral hospitals to obtain adequate drugs when compared to those patients those obtained adequate drugs from the proximal facilities (AOR of 95% CI=2.366(1.013-5.526).

This finding is in line with the finding of the study conducted in Tanzania shows that reasons for patients referral from lower level health facilities were lack of expertise and equipment were the most common factors given for self-referral (96.3%) and about half of the lower level health facilities reported lack of drugs (53.8%) and due to the factors patients bypassed the proximal health facilities [12].

From the study result, 65.8% of the self-referred outpatients perceived the quality of health services at proximal health facility was poor and preferred self-referral to referral hospitals. Consequently, the odd of self-referral patients perceived quality of health services at proximal health facility was poor were almost three times more likely to self-refer than those patients perceived the quality of health services at proximal health facilities was good (AOR of 95% CI=2.996(1.418-6.328).

Similarly the study conducted in Kenya identified that quality of health services is one of institutional determinant of patients’ self-referral to referral health facilities [17]. In addition to this according the study conducted in Chad patients those know the other health care providers do not offer the specific services quality they need 13.3% higher probability the patients had bypassed the proximal providers to seek cares [18].

The patients those perceived the location of the proximal health facility was inconvenient and not accessible to transportation were almost four times more likely to self-refer compared to those patients perceived the location of the proximal health facility was convenient and accessible to transportation (AOR of 95% CI=3.750(1.560-9.017) and CI=3.879(1.651-9.114) respectively.

Similarly the study conducted in Kenya revealed that location of health facilities is one of the institutional determinants cited as patients as the reasons why they prefer to seek health services from referral health facilities bypassing lower level health facilities [17] and study in Honduras revealed that due to geographical accessibility 84% of 111 cases patients directly referred to higher referral hospitals by passing health facilities that functioning as intermediate level between lower level health centers and higher referral health facilities [18]. This might be due to when the health facility is at convenience location and access to transportation patients easily travel to and visit the facility to get health services.

## 5. Conclusion

The proportion of outpatient self-referral to referral hospitals bypassing the proximal health facilities was high. Referral information on referral linkage of health facilities, perception of patients about their severity of illnesses and quality of health services as well as confidence of patients to get the type of health care provider they want to see at the facilities were the individual factors that influence outpatient self-referral to referral hospitals where as unavailability of laboratory services, adequate drugs that patients expect to get from the health facilities, convenience location and accessibility of the health facilities to transportation were major institutional factors that influence the outpatients self-referral to referral health facilities.

We strongly suggest that the Federal Ministry of Health and Regional Health Bureau should develop monitoring systems of referral linkage of health facilities to strengthen referral systems at all level of health facilities and ensure health services needed to be delivered at lower level health facilities. The health care providers should create awareness in the community to visit the proximal health facilities early when they sick for their any health problems and about referral linkages of health facilities. Further study research may be useful in to evaluate the costs of outpatient self-referral with the perspective of patients and health care system.

## Acknowledgement

We are grateful to the study participants and our special thanks go to Jimma University for its financial support of the study.

## References

1. Henery J .Focus on health care disparities key facts. Kaiser Family Foundation.2012; 2–3.

2. Syed M, Bushra B, Iqbal A, etal. Bangladesh Health system Review. Health systems in Transitions. WHO, 2015; 5(3): 142–152.

3. Macintyre K, Megan L, David R, etal. Barriers to Referral in Swaziland: Perceptions from providers and Clients of System under Stress. World and medical and health policy. PSO 2011.Manuscript 1183; 24–26.

4. Health Sector Transformation plan /HSTP2015/2016-2020/. Ethiopian federal ministry of health. Addis Ababa, Ethiopia. 2015; 41–43.

5. Studying the healthcare network and Analyzing Disrupting Health sector. World health organization. WHO/HAC/MAN/.2008; 8 rev1:281–289.

6. Mojaki ME, Basu D, Letskokgohka ME, Govender M. Referral steps in district health system are side-stepped. Scientific letters. SAMJ. 2011; 101(2): 109.

7. Abrahim O, Linnander E, Mohammed H, Fetene N, Bradley E. A Patient-Centered Understanding of the Referral System in Ethiopian Primary Health Care Unit.PloSONE.2015; 10(10):1–10. DOI:10.1371/journal.pone.0139024

8. Wolkite O, Waju B, GebeyehuT. Magnitude and Determinants of Self-Referral of Patients at a General Hospital, Western Ethiopia. Science Journal of Clinical Medicine. 2015; 4(5): 86–92. Doi:10.11648/j.sjcm.20150405.12

9. Stewart G. The Increasing Importance of Physician to Physician Referrals. Health care success. 2014.

10. Karimollah H. Sample size determination in epidemiological studies. Caspian J Intern Med. 2011;2(4): 289–298.

11. Nath B, Kumari R, Tanu N. Utilization of the health care delivery system in a district of north India. East African Journal of Public Health.2008; 5(3):147–153.

12. Daudi O, Naboth A, Lawrence M, Leonard L. Referral pattern of patients received at the national referral hospital: Challenges in low income countries. East Africa Journal of Public Health.2008; 5(1):7–8.

13. Yaffee A, Whiteside L, Oteng R, etal. By passing proximal health care facilities for acute care: a survey of patients in a Ghanaian Accident and Emergency Centre. Tropical medicine and International Health. 2012; 17(6):777–781. Doi:10.1111/j.1365-3156.2012.02984

14. Emma S, Jonathon K. Direct access and patient self-referral. WCPT Keynotes/ Direct access. World Confederation for Physical Therapy. 2011; 1–2.

15. Teresita A. The determinants of health care seeking and bypassing of health care facilities in Kenya. 2014; 19–24.

16. Christopher B, Jonathan P, Jinnet F, etal. Self-referral in Point-of-Service Health Plans.JAMA.2010; 285(17):2223–2234.

17. Mahindra F. Determinants of self directed referral amongst patients seeking health services at Kenyatta national hospital. Nairobi. 2013; 39–50.

18. Kumiko O, Victor M, Naruo U, Gen O. Study of a patient referral system in the Republic of Honduras. Health Policy and Planning. Oxford University Press.1998; 13(4):436–440.

